# Lower-limb express visuomotor responses are spared in Parkinson’s Disease during step initiation from a stable position

**DOI:** 10.1101/2024.11.29.625631

**Authors:** Lucas S. Billen, Jorik Nonnekes, Brian D. Corneil, Vivian Weerdesteyn

## Abstract

While motor impairments have been extensively studied in Parkinson’s Disease, rapid visuomotor transformations for flexible interaction with the environment have received surprisingly little attention. In recent years, such rapid visuomotor transformations have been studied in the form of express visuomotor responses (EVRs), which are goal-directed bursts of muscle activity that are thought to originate from superior colliculus, reaching the periphery via the tecto-reticulospinal pathway.

Here, we examined EVRs in the lower limbs during goal-directed step initiation in 20 people with Parkison’s Disease (PwPD) and 20 age-matched healthy control participants (HC). As lower-limb EVRs in the young have been shown to interact with postural control - which is often affected in PwPD – we manipulated the postural demands by varying initial stance width and target location. In the low postural demand condition, EVRs were robustly present in both the PwPD (17/20) and HC (16/20) group. However, when postural demands were high, EVRs were largely absent in both groups and, instead, strong anticipatory postural adjustments (APAs) were required prior to foot off. EVR magnitudes were, on average, stronger in PwPD compared to HC, but they decreased with increasing disease severity, suggesting that the EVR network may become compromised or down-regulated in later stages of the disease. While APA magnitudes were smaller in PwPD compared to HC, subsequent stepping performance (step reaction time, duration, size, velocity) was remarkably similar between the two groups. We suggest that the EVR network may be upregulated in the early stages of Parkinson’s disease in order to compensate for some of the emerging motor deficits experienced in daily life.

## Introduction

Parkinson’s disease (PD) results in both motor and non-motor symptoms. Gait and balance impairments are a hallmark of the disease (Caetano et al., 2018; Clarke, 2007; Contreras & Grandas, 2012; Palakurthi & Burugupally, 2019). Despite extensive research into gait and balance impairments in PD, the ability for people with PD (PwPD) to rapidly and flexibly change stepping behavior in response to changes in an inherently dynamic environment has received relatively little attention.

This ability is essential in everyday life, for example to ensure safe locomotion on uneven terrain, as it involves the complex interplay between movement adjustments of the stepping leg and postural control. The few studies that have looked at this suggest that this ability is impaired in people with PD, in parallel with a potentially higher risk of falling (Borm et al., 2024; Caetano et al., 2018; Geerse et al., 2018). Yet, the underlying mechanisms have remained elusive.

To facilitate such rapid goal-directed stepping behavior, involvement of a reflexive, yet highly adaptive, fast visuomotor network has been proposed (Queralt et al., 2008; Reynolds & Day, 2005, 2007). In the upper limb and neck, these rapid visuomotor transformations are thought to originate in the midbrain superior colliculus from where they are relayed to the brainstem reticular formation and subsequently to the motor periphery via the tectoreticulospinal tract (Boehnke & Munoz, 2008; Corneil et al., 2004; Corneil & Munoz, 2014). Indeed, a network involving the superior colliculus has been proposed to underlie the initiation of our most rapid, visually-guided actions, not only for rapid oculomotor movements such as express saccades (Munoz et al., 2000), or orienting head movements (Corneil et al., 2004; Rezvani & Corneil, 2008), but also for mid-flight adjustments of either the upper (Day & Brown, 2001) or lower (Fautrelle et al., 2010; Weerdesteyn et al., 2004) limbs. In recent years, bursts of short-latency muscle activity occuring in a time-locked window ∼100ms after appearance of a salient visual stimulus (termed express visuomotor responses, EVRs), have been proposed to arise from signaling along the tectoreticulospinal tract (Corneil et al., 2004; Pruszynski et al., 2010). In the upper limb, EVRs can be generated from either a stable starting posture or during mid-flight reaching adjustments (Kozak et al., 2019), are directionally tuned to the location of the stimulus (Contemori et al., 2023; Gu et al., 2016) and they facilitate the rapid goal-directed movement towards the target, as stronger EVRs correlate with faster subsequent reaction times (Gu et al., 2016; Pruszynski et al., 2010; Wood et al., 2015).

Could degradation of the fast visuomotor network underlie deficits in visually-guided stepping in PD? Neuropathologically, the tectoreticulospinal pathway may be directly affected, as there is early-stage degeneration in the brainstem, which spreads to higher-order areas as the disease progresses (Braak et al., 2003; Diederich et al., 2014; Jubault et al., 2009; Seidel et al., 2015). Furthermore, disease-mediated changes in the inputs to the superior colliculus, for example from the basal ganglia or pedunculopontine nucleus, may lead to over- or under-excitability of the fast visuomotor network depending on the excitatory or inhibitory nature of the projections involved. For the upper limb, previous reports have reached differing conclusions on whether PD impacts fast corrections of reaching movements in mid-flight, as one study reported deficits (Desmurget et al., 2004) whereas another reported that this ability was retained (Merritt et al., 2017); the initiation of interceptive movements of the upper limb was also reported recently to be spared (Fooken et al., 2022). To date, the only study of upper limb EVRs in PD reported that they were spared in PD (Gilchrist et al., 2024). Importantly, disease severity differed between the studies, with the PD cohort in the Desmurget study being the most severely affected (average UPDRS Motor subscale score of 36.41, OFF state) and the cohort in the Merritt study (average UPDRS Motor subscale score of 11.07, OFF state) being least affected. This discrepancy suggests a potential effect of disease severity on the integrity of the fast visuomotor network.

Here, we primarily aimed to examine EVRs in the lower limbs in PD. This is of particular interest because of the recently demonstrated interplay between lower limb EVRs and postural control (Billen et al., 2023). Compared to reaching movements, rapid stepping responses are more posturally-demanding, usually requiring strong anticipatory postural adjustments (APAs) prior to step initiation that shift the centre of mass towards the stance side. In our study (Billen et al., 2023), we found a reciprocal relationship where stance-side EVRs consistently preceded and contrasted the subsequent step-side APAs: EVRs were robustly expressed when stepping in a low-postural demand condition that did not require APAs, but EVRs were suppressed when stepping from in a high postural demand condition requiring APAs. The downregulation of EVRs in the high postural demand condition may reflect a prioritization of balance over speed during step initiation. Importantly, in this context, the occasional erroneous expression of stance-side EVRs negatively impacted task performance, as EVR expression was followed by larger compensatory (stepping-side initiated) APAs prior to step onset and consequent delays in step reaction times (Billen et al., 2023).

These observations, which were made in young healthy adults, raise the question about what happens in PD, and in aging. APAs are substantially impaired in PD (Halliday et al., 1998; Hass et al., 2005; Lin et al., 2016). If, in addition to impaired APA expression, the posturally-dependent regulation of EVRs is also affected in PD, this may be reflected in balance demands not being met, increasing the risk of falling. Here, we aim to better understand this complex interaction between EVRs and APAs in PD, and in aging. Like the paradigm used in a previous study (Billen et al., 2023), participants performed a visually guided stepping task while we manipulated the postural demands of the upcoming step.

Behaviorally, we expect APAs and stepping behavior to be significantly impaired in PwPD compared to an age-matched healthy control group. Based on the more recent findings of intact upper-limb EVRs in PD, we hypothesize that the EVR network is inherently spared, at least in those with mild to moderate disease severity. Further, to examine potential degradation of the fast visuomotor network in more advanced disease stages, we determined the relationship between EVR expression and disease severity (as measured with the MDS-UPDRS part III) within the PwPD.

## Materials and Methods

### Participants

Twenty participants with idiopathic Parkinson’s Disease (12 males, 8 females, 65.8±7.1 years) and twenty age-matched healthy older adults (12 males, 8 females, 66.5±8 years) participated in this study. Inclusion criteria involved a BMI under 25 kg/m^2^ to minimize the coverage of muscles by adipose tissue, which could compromise the quality of surface EMG recordings. Exclusion criteria were any (additional) visual, neurological, or motor-related disorders that could influence the participant’s performance in the study. PwPD were not required to deviate from their medication schedule. Before the start of the experiment, each PwPD participant completed the MDS-UPDRS assessment. The average total MDS-UPDRS score was 42.4 (*SD* = 16.6) with an average part III (motor subscale) score of 28.9 (*SD* = 13.2, min: 12, max = 52). None of the PwPD regularly experienced freezing of gait, so the New Freezing of Gait Questionnaire (N-FOG; Nieuwboer et al., 2009) was not administered. The study protocol was reviewed by the medical ethics committee (CMO Arnhem-Nijmegen, 2022-16109) and conducted in accordance with the latest version of the Declaration of Helsinki. All participants provided written informed consent prior to participation and were free to withdraw from the study at any time.

### Data collection & experimental design

The experiment was performed using a Gait Real-time Analysis Interactive Lab (GRAIL, Motek Medical, The Netherlands), as previously described in Billen et al. (2023). In short, the experimental setup included an M-gait dual-belt treadmill with two embedded force plates (sampled at 2000 Hz, GRAIL, Motek Medical, The Netherlands) to measure ground reaction forces, a 10-camera 3D motion analysis system (sampled at 100 Hz, Vicon Motion Systems, United Kingdom) and a projector (Optoma, UK) to project all visual stimuli. Muscle activity of gluteus medius was recorded using Ag/AgCl surface electrodes and a Wave Wireless electromyography system (sampled at 2000 Hz, Wave Wireless EMG, Cometa, Italy). GM was chosen instead of tibialis anterior (TA; a muscle commonly reported as being involved in APAs), because our previous study showed that the initial recruitment of TA did not differ after left or right target presentation. Electrodes were placed in accordance with the SENIAM guidelines (Hermens & Merletti, 1999) and signal quality was checked prior to the experimental task. Trials were started manually via the D-flow software (Motek Medical, The Netherlands) by the experimenter. A secondary peripheral target measured by a photodiode (TSL250R-LF, TAOS, USA) was used to account for small variable delays in target presentation. All reported measures (i.e. EMG and force plate measures) were aligned to the moment of stimulus presentation detected by the photodiode.

Participants stood on the stationary M-Gait with each foot placed on a separate force plate. They performed a modified version of an emerging target paradigm (Kozak et al., 2020) known to promote EVRs (Contemori et al., 2021a; Kozak & Corneil, 2021), which we modified for a stepping task (Billen et al., 2023). The initial stance position was indicated by the projection of small circles at the desired foot location. The stepping task was projected on the treadmill in front of the participant (Figure 1). Each trial started with the appearance of a projected stationary visual target in front of the participant (130cm from participant). The target started moving towards the participant with a constant velocity, then it disappeared behind an occluder (a light blue rectangle) for a fixed interval of 750ms and subsequently it reappeared randomly as a single flash (48ms, i.e. 3 frames) in front of the left or right foot of the participant. Participants were instructed to perform a full stepping movement upon reappearance of the target, using the leg on the side of target appearance (i.e. step with the left leg when the target appeared on the left side and vice versa for the right leg) and placing the stance leg next to the stepping leg in order to complete the stepping movement. After completing the trial, the participant returned to the starting position, and the subsequent trial began. As was done previously (Billen et al., 2023), we instructed participants to initiate and complete the step as rapidly as possible. As a slight amendment to the previous instructions, we also aimed to further increase the participant’s motivation to step fast by instructing them to imagine that the reappearing target was a small flame that they need to extinguish as rapidly as possible by stepping onto it. This was done following pilot experiments showing that older individuals were less inclined to step as fast at the younger cohort in our previous. Frequent reminders were also provided throughout the experiment.

**Figure 1.**
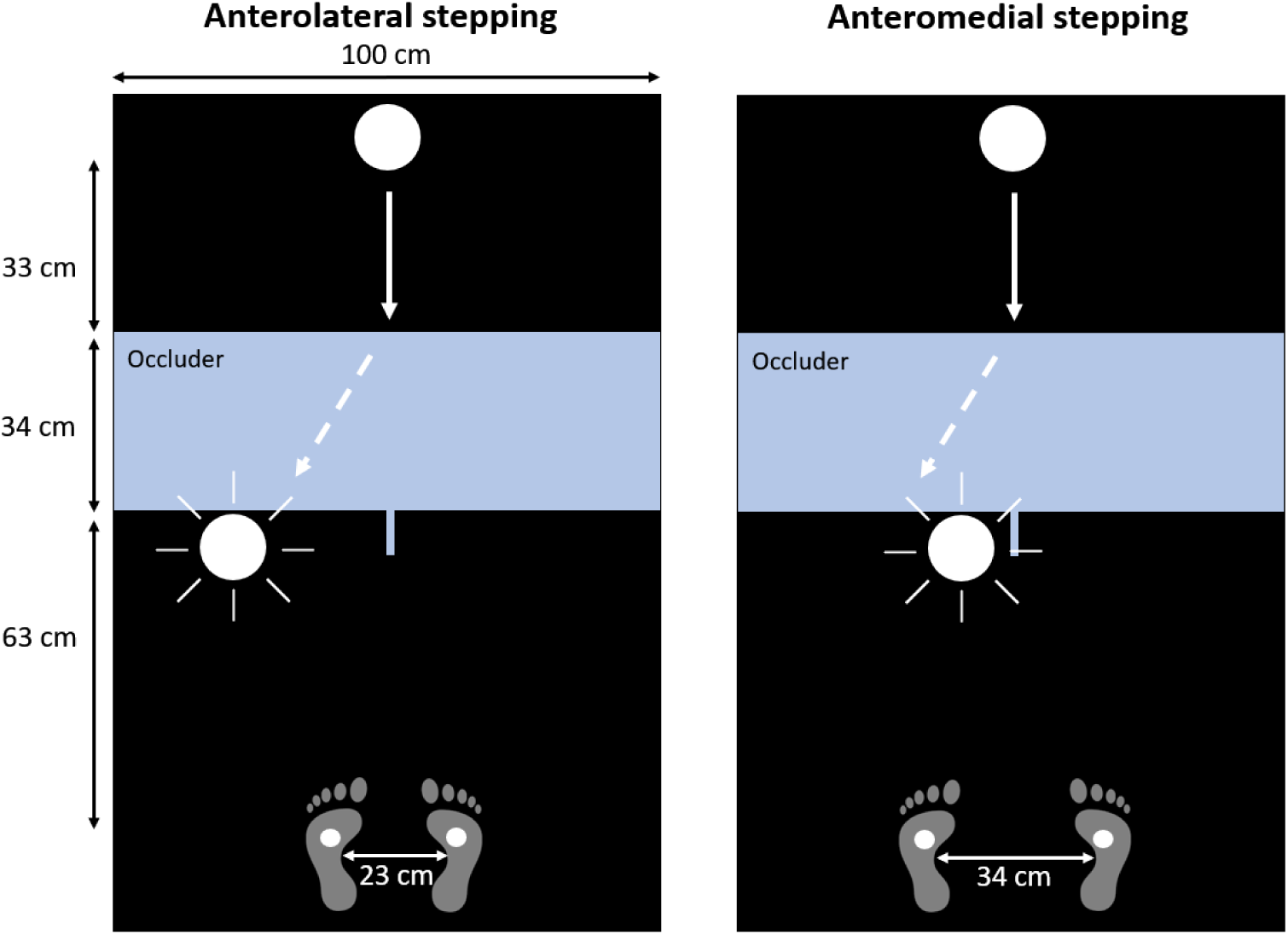
Experimental setup of the emerging target paradigm. The paradigm was projected on the floor in front of the participants. Participants placed their feet on two projected dots. The visual target moved down towards the participants, disappeared behind the occluder, and then, in this example, reappeared in front of left foot of the participant. Participant stepped onto the target upon reappearance, requiring either an anterolateral (left figure) or anteromedial (right figure) stepping response

In separate blocks of trials, the postural demands of the upcoming step were manipulated by presenting the stepping target either anterolaterally (stepping forward and outward from a narrow stance) or anteromedially (stepping forward and inward from a wide stance) in front of the stepping foot. Altering the target location and stance width dynamically modifies the postural demands of the stepping task. Stepping medially from a wide stance increases balance demands and, as a result, requires an anticipatory postural adjustment (APA). Conversely, stepping laterally from a narrow stance toward anterolateral targets reduces these demands. Participants completed 4 blocks of 75 trials (300 in total). Each block consisted of either only anterolateral targets or anteromedial targets and the order of the blocks was counterbalanced. Participants were informed about the condition before each block. Target side (left/right) was randomized on each trial.

### Data processing and analysis

Incorrect trials were excluded from the analysis and were defined as trials in which participants stepped towards the wrong direction or initiated a stepping movement with the contralateral foot. Data analysis was performed using custom-written MATLAB scripts (version 2019a).

#### Reaction time

Stepping reaction time (RT) was defined as the interval between the appearance of the visual target, measured using a photodiode, and the moment the stepping foot was lifted off the ground.

Consistent with previous studies, foot-off was identified as the first instance where the vertical ground reaction force (Fz) dropped below one percent of the participant’s body weight (Rajachandrakumar et al., 2017).

#### EVR presence and latency

Raw EMG signals were first band-pass filtered between 20 and 450 Hz and subsequently rectified and low-passed filtered at 150 Hz with second-order Butterworth filters. To determine the presence and latency of lower limb EVRs, we used a time-series receiver-operating characteristic (ROC) analysis, as described previously (Billen et al., 2023). Briefly, the target side (left vs. right) and postural condition (anterolateral vs. anteromedial) were used to group the EMG data. EMG activity was then compared between leftward and rightward steps within either condition. An ROC analysis was carried out, which, for each sample between 100 ms prior to and 500 ms following the visual stimulus appearance, computed the area under the ROC curve (AUC). This measure shows the likelihood that an ideal observer, relying just on EMG activity, could distinguish between the two sides of stimulus presentation. The AUC value range is 0 to 1, where 0.5 denotes chance discrimination and 1 or 0 denotes correct or incorrect discrimination, respectively. We determined the discrimination threshold to be 0.6 in accordance with earlier studies (Gu et al., 2016). Within the pre-specified EVR epoch of 100-140 ms following stimulus presentation, the time of earliest discrimination was determined as the moment at which the AUC exceeded the discrimination threshold and stayed above the threshold for 16 out of 20 consecutive samples.

#### Response magnitude in EVR window

The response magnitude in the EVR window (in this paper, synonymous to the term “EVR magnitude”) was calculated for each condition within each participant, regardless of whether an EVR was detected. On a single trial basis, the mean EMG activity of the 20ms window centered around the maximum EMG activity during the EVR epoch (100-140 ms) was calculated. Magnitudes were then normalized against the median peak EMG activity (in the interval from 140 ms to foot-off) during anterolateral stepping of the respective participant. EMG magnitudes of all trials were then averaged per condition.

#### APA onset and magnitudes

As with EVRs, the onset of an APA was determined using a time-series receiver operating characteristic (ROC) analysis on EMG data of gluteus medius to determine the timepoint at which stepping-side GM activity increased significantly compared to stance-side activity, signifying APA initiation. The discrimination threshold was set to 0.6 (this threshold had to be crossed for 8 out of 10 consecutive trials) and the ROC analysis was carried out in the time window of 100-300ms following target reappearance.

APA magnitude was defined based on the mean ground reaction forces. In the interval from 140ms after target appearance (i.e., the end of the EVR window) and foot-off, the maximum vertical ground reaction force component (Fz) underneath the stepping leg was determined and corrected for baseline. Subsequently, the difference between this maximum and its corresponding ground reaction force underneath the stance leg was calculated and then normalized to percent total body weight (%BW).

## Statistical analysis

Statistical analyses were performed using MATLAB (version R2019a). The level of significance was set to *p* < .05 for all analyses. Repeated Measures ANOVAs were performed to study whether EVR magnitudes, APA magnitudes as well as stepping parameters (stepping RT, velocity, size, duration) differed between *postural demand* (anterolateral/anteromedial stepping) and between *groups* (PD/HC).

To compare EVR prevalence between the HC and PD groups we used Fisher’s exact test. Two-sample t-tests were used to test whether APA onset times during anteromedial stepping and EVR latencies during anterolateral stepping differed between the PD and HC groups; and whether UPDRS scores differed between PwPD with and without EVR expression. Spearman’s rank correlation coefficients were determined to study whether APA and EVR magnitudes were associated with UPDRS scores.

## Results

Any differences between stepping sides (left/right) in behavioral outcomes and EMG-related outcomes were not significant. Within the PD group, differences between the more and the less affected leg were also not significant. We therefore averaged all outcomes across sides.

## Error rates

All participants completed the task with low error rates. Error rates were significantly lower during anterolateral stepping compared to anteromedial stepping in both the HC group (anterolateral: 1.9%, anteromedial: 8.4%, t(19) = -5.62, *p* < .001) and the PD group (anterolateral: 4.2%, anteromedial: 9.1%, t(19) = -3.54, *p* = .002). Differences between groups were non-significant, but error rates differed greatly between individuals. For example, in the PD group, the most error-prone participant made 26 errors in the anteromedial stepping blocks (17.3% of trials) and 14 in anterolateral stepping (9.3%), whereas others made virtually no errors. Similarly, in the HC group, the most error-prone participant made 18 errors in anteromedial stepping (12%) and 6 errors during anterolateral stepping (4%).

## Apart from APAs, behavioral outcomes were unaffected in PD

We found a significant main effect of *postural demand* (F(1) = 597.0, *p* < .001) on APA magnitudes, with APAs in anterolateral stepping being either completely absent or small in magnitude (HC group, *M* = .08 %BW, *SD* = 0.1; PD group, *M* = .06 %BW, *SD* = 0.1) but strongly expressed in anteromedial stepping (HC group, *M* = .55 %BW, *SD* = 0.12; PD group, *M* = .41 %BW, *SD* = 0.19). APAs were on average smaller in the PD than the HC group (*group,* F(1) = 9.7, *p* < .01; see Figure 2A). In the anteromedial stepping condition, APA onset times did not differ significantly between the HC group (*M* = 162ms, *SD* = 17ms) and the PD group (*M* = 160ms, *SD* = 18ms; *T*(38) = - 0.20, *p* = .84).

**Figure 2.**
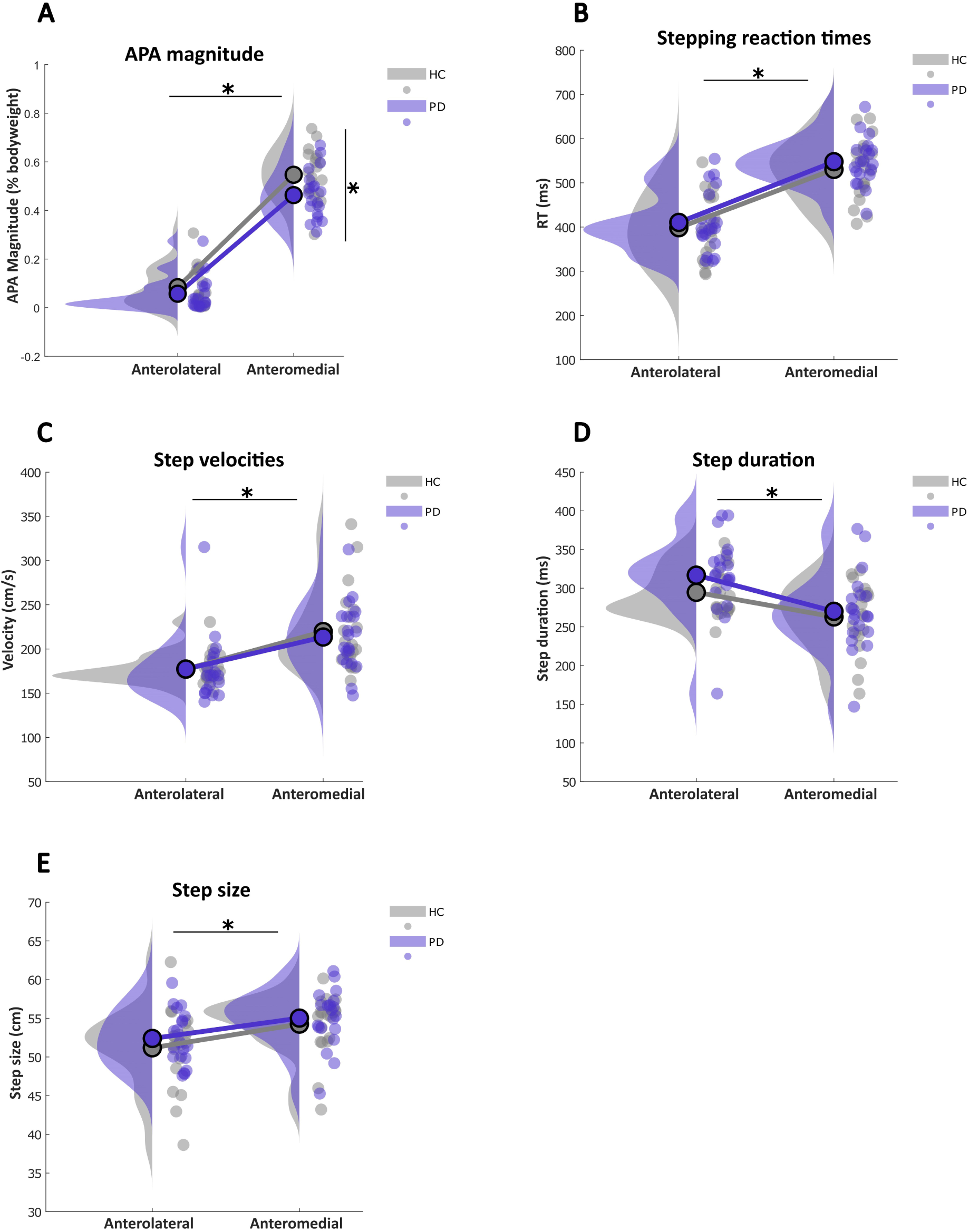
**A-E.** Behavioral outcomes of all participants from the HC (purple) and PD (grey) group for anterolateral and anteromedial stepping. The dots indicate individual averages, averaged across left and right steps. The density plots indicate the distribution of the data. The asterisks indicate significant differences (p < .05) between postural demands (horizontal) and groups (vertical)

Stepping reaction times were significantly faster for anterolateral steps (HC: *M* = 399ms, *SD* = 76ms; PD: *M* = 414ms, *SD* = 75ms) than for anteromedial steps (HC: *M* = 530ms, *SD* = 71ms; PD: *M* = 527ms, *SD* = 80ms; F(1) = 148.3, *p* < .001). Step RTs did not differ between groups (*p* = .19; see Figure 2B).

To investigate whether the stepping behavior itself was affected in PwPD, we investigated step velocity, size, and duration. There were significant effects of *postural demand* on all of these measures, with the steps in the anteromedial condition resulting in significantly higher step velocities (anterolateral: *M_HC_* = 177 cm/s, *SD_HC_*= 20 cm/s; *M_PD_* = 176 cm/s, *SD_PD_* = 42 cm/s; anteromedial: *M_HC_* = 220 cm/s, *SD_HC_* = 50 cm/s; *M_PD_* = 210 cm/s, *SD_PD_* = 41 cm/s; *F*(1) = 39.42, *p* < .001), shorter step durations (anterolateral: *M_HC_* = 295 ms, *SD_HC_* = 34 ms; *M_PD_* = 316 ms, *SD_PD_* = 58 ms; anteromedial: *M_HC_* = 263 ms, *SD_HC_* = 45 ms; *M_PD_* = 277 ms, *SD_PD_* = 55 ms; *F*(1) = 26.91, *p* < .001) and slightly larger step sizes (anterolateral: *M_HC_* = 51 cm, *SD_HC_* = 6 cm; *M_PD_* = 52 cm, *SD_PD_* = 3 cm; anteromedial: *M_HC_* = 54 cm, *SD_HC_* = 4 cm; *M_PD_* = 55 cm, *SD_PD_* = 4 cm; *F*(1) = 17.67, *p* < .001). There were no significant group differences in any of these outcomes (*p* > .05; see Figure 2C-E).

## EMG and force plate data in a representative PD and control participant

The upper half of Figure 3 shows the data of a representative participant from the control group. The first column depicts the mean EMG activity across trials of left GM when it is on the stance side (dark blue) and when it is on the stepping side (light blue). On the stance side in the anterolateral condition (top row of Fig 3), there is an initial burst of muscle activity at ∼100ms. This is the Express Visuomotor Response, as also demonstrated by the time-series ROC plot shown above the EMG trajectories. In the trial-by-trial data (colored heatmaps in second column), the EVR is visible as a vertical band of activity that is independent of the subsequent reaction time (EVR) and time-locked to target presentation (red rectangle on stance-leg heatmap). Following the EVR, there is a second burst of activity, which corresponds to the voluntary muscle activity related to step initiation. On the fastest trials, this activity fuses with the EVR, but it is otherwise separate from it. Activity on GM in the stepping leg remains relatively low throughout the initial phase of step initiation, and only ramps up around foot-off. The force plate data (column 4) shows an immediate increase of vertical force on the stance side and decrease on the stepping side, indicating that APAs were generally not executed.

**Figure 3.**
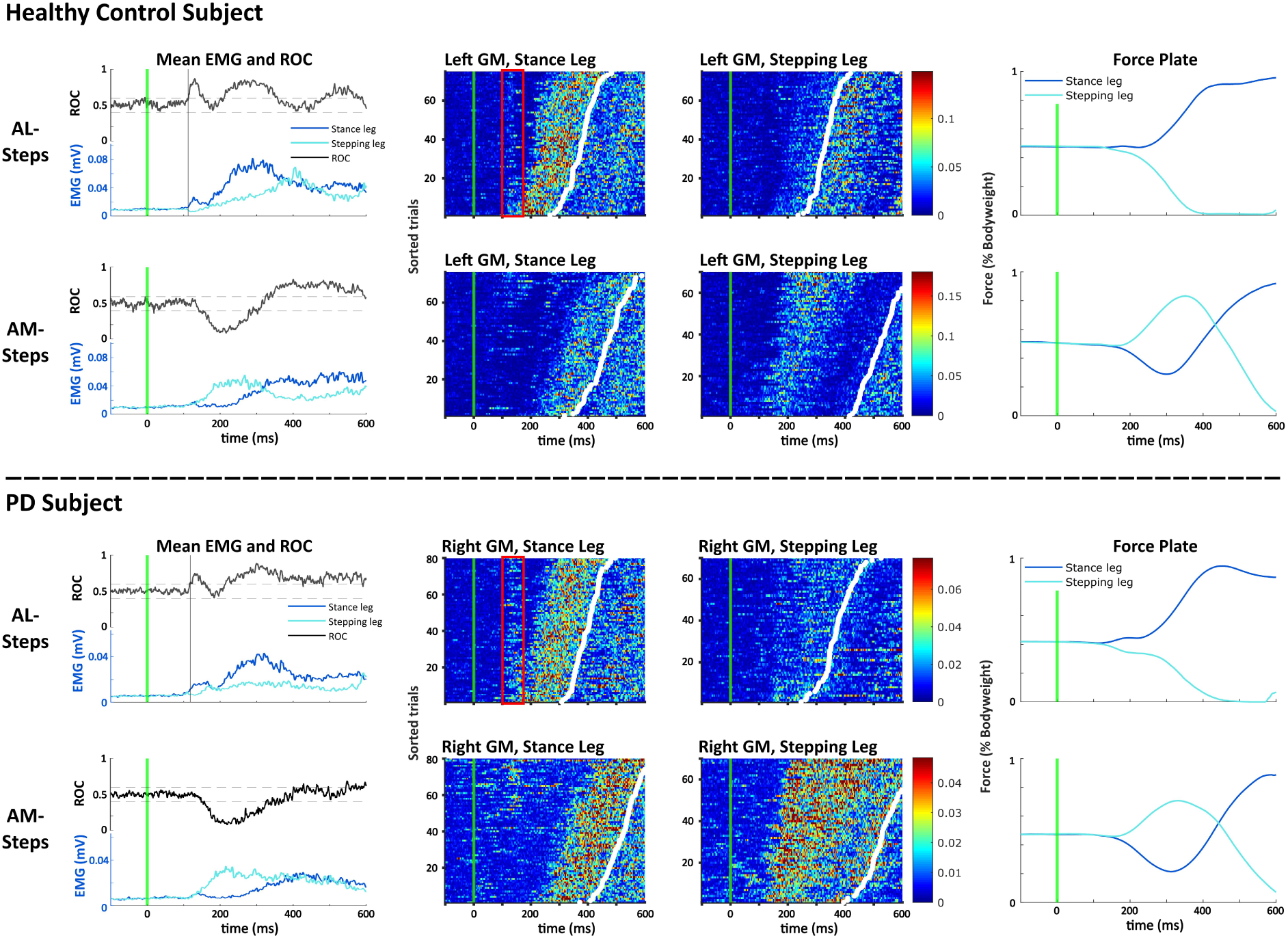
GM muscle activity, time-series ROC analysis and force plate data of an exemplar participants from the HC (top half, left GM) and PD (bottom half, right GM) group. Data is separated based on stepping condition (AL = anterolateral; AM = anteromedial). Each condition is presented on a separate row. All data are aligned to visual stimulus onset (green line). **Column 1:** shows mean EMG activity for the stance-side (dark blue) and the stepping side (light blue). The time-series ROC curve is shown in black. Discrimination times within the EVR epoch (100-140 ms) are indicated by the black vertical line. **Columns 2 and 3:** Trial-by-trial EMG activity of left GM when on the stance side (column 2) or the stepping side (column 3). Intensity of color conveys the magnitude of EMG activity. Each row represents a different trial. Trials are sorted by RT (white dots). **Column 4:** Mean vertical force (Fz) exerted by the stance (dark blue) and stepping leg (light blue). The initial increase in force under the stepping leg in the anteromedial condition corresponds to the APA.

In anteromedial stepping (second row), the general patterns of muscle activity look quite different. As is visible on the average EMG traces, there is a slight increase on stance side GM right around the EVR-window, which, looking at the trial-by-trial activity, is caused by a few trials showing EVR activity during trials with slow step RTs. This initial burst of activity is promptly suppressed and the stance side only becomes active again shortly prior to foot-off. Instead, and in contrast to the anterolateral condition, stepping-side GM shows a pronounced increase in muscle activity around 150ms after target reappearance which corresponds to strong APA expression. This is visible as an increase in vertical force at the stepping side at ∼160ms (column 4) which induces a CoM shift towards the stance side.

The lower half of Figure 3 depicts data of a representative participant with PD. Overall, the pattern of muscle recruitment, the reaction times and the ground reaction forces are remarkably similar to the control participant. In anterolateral stepping, strong and robust EVRs were evoked on most trials, which is visible in both the average EMG traces as an increase in stance leg activity (first column, first row), as well as on the trial-by-trial plot (second column, red rectangle) as a vertical band of time-locked activity around 120ms. Similar to the control participant, stepping side GM remains relatively silent during anterolateral stepping, resulting in the absence of APAs.

In contrast to anterolateral stepping, APA-related activity in this representative participant with PD is again very strong during anteromedial stepping. Stepping-side GM becomes active at around 140ms whereas stance-side GM remains relatively silent, which ‘push-off’ activity induces the ensuing CoM shift towards the stance side, as shown by the force plate data. EVRs are absent in most trials in this participant during anteromedial stepping, but, similar to the HC participant, there are hints of EVR expression on trials with slower step RTs.

## Are EVRs spared in PD?

To establish EVR expression in both groups, we first investigated EVR prevalence across groups and postural conditions. During anterolateral stepping, EVRs were robustly present in the majority of participants in both groups (HC: 16/20, 10 with bilateral EVRs; PD: 17/20, 12 with bilateral EVRs; *p* = .68). Average EVR latencies were similar across both groups (HC: *M* = 120ms, *SD* = 7.5; PD: *M* = 119ms, *SD* = 9.3; *p* > .75; see Figure 4A). In anteromedial stepping, none of the participants of either group exhibited EVRs.

**Figure 4.**
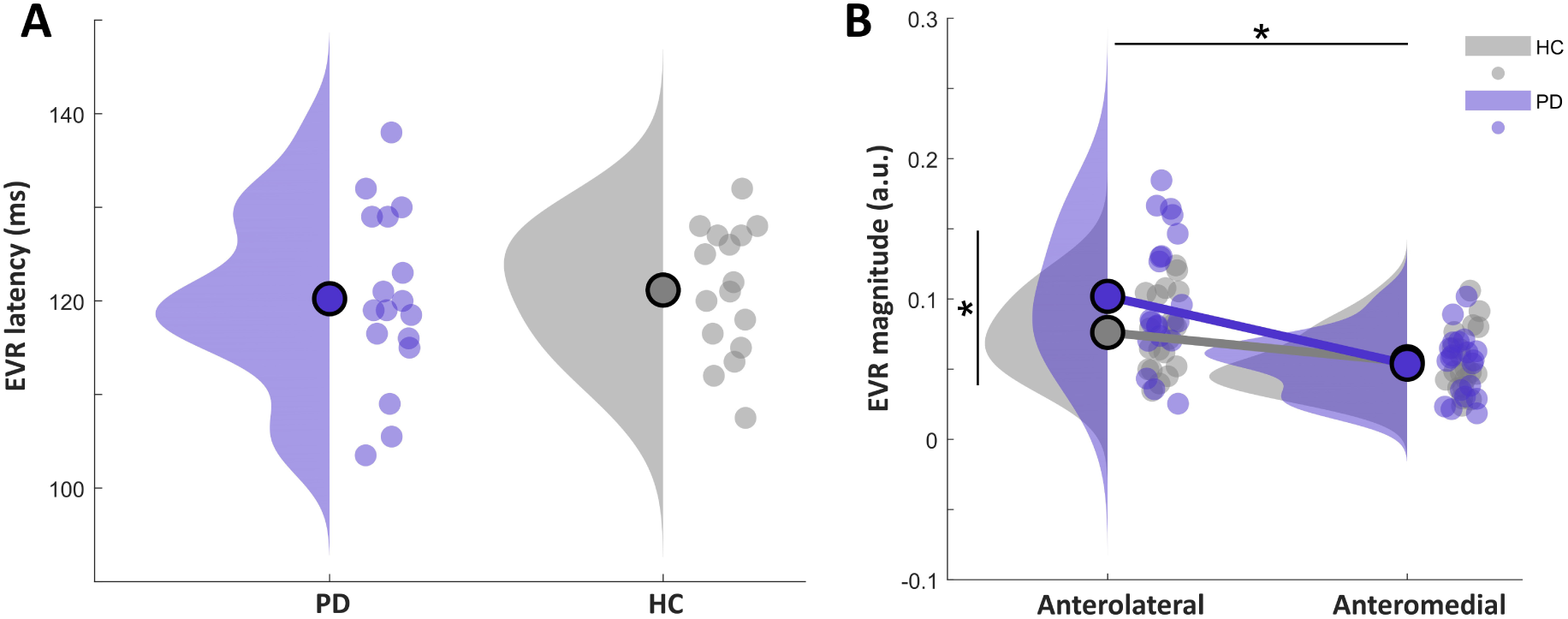
**A:** Average EVR latencies of all participants in the PD (purple) and the HC (grey) group during anterolateral stepping. If participants had bilateral EVRs, the average latency across sides are displayed here. Participants without EVRs on either side are not displayed here. **B:** Response magnitudes within the EVR window of all participants from the HC (grey) and PD (purple) group for anterolateral and anteromedial stepping. The dots indicate individual averages, averaged across left and right steps. The density plots indicate the distribution of the data. The asterisks indicate significant differences (p < .05) between postural demands (horizontal) and groups (vertical). There was also a significant interaction between groups and postural demand.

Investigating the response magnitude within the EVR window, we found significant effects of *group* (F(1) = 4.1, *p* = .045) and *postural demand* (F(1) = 33.7, *p* < .001), as well as a significant *group x postural demand* interaction (F(1) = 5.3, *p* = .022). In anterolateral stepping, response magnitudes were higher in the PD group (*M* = .10 a.u., *SD* = 0.06) compared to the HC group (*M* = .08 a.u., *SD* = 0.03), whereas response magnitudes were similarly small across both groups in the anteromedial stepping condition (HC: *M* = .06 a.u., *SD* = 0.03; PD: *M* = .06 a.u., *SD* = 0.03; Figure 4B).

We further evaluated whether stronger EVRs preceded faster stepping reaction times during anterolateral stepping, as previously reported in reaching (Pruszynski et al., 2010) and stepping (Billen et al., 2023). Indeed, on a trial-by-trial basis, there were negative, albeit non-significant, correlations in both the HC group (*ρ* = -.28, *p* = .21) and the PD group (*ρ* = -.41, *p* = .054), indicating that higher EVR magnitudes tended to precede shorter stepping reaction times. On a group-level (average EVR magnitude and step RT per participant), we observed significant negative correlations in the PD group (*ρ* = -.72, *p* < .001), but not the HC group (HC: *ρ* = -.42, *p* = .066, Figure 5).

**Figure 5.**
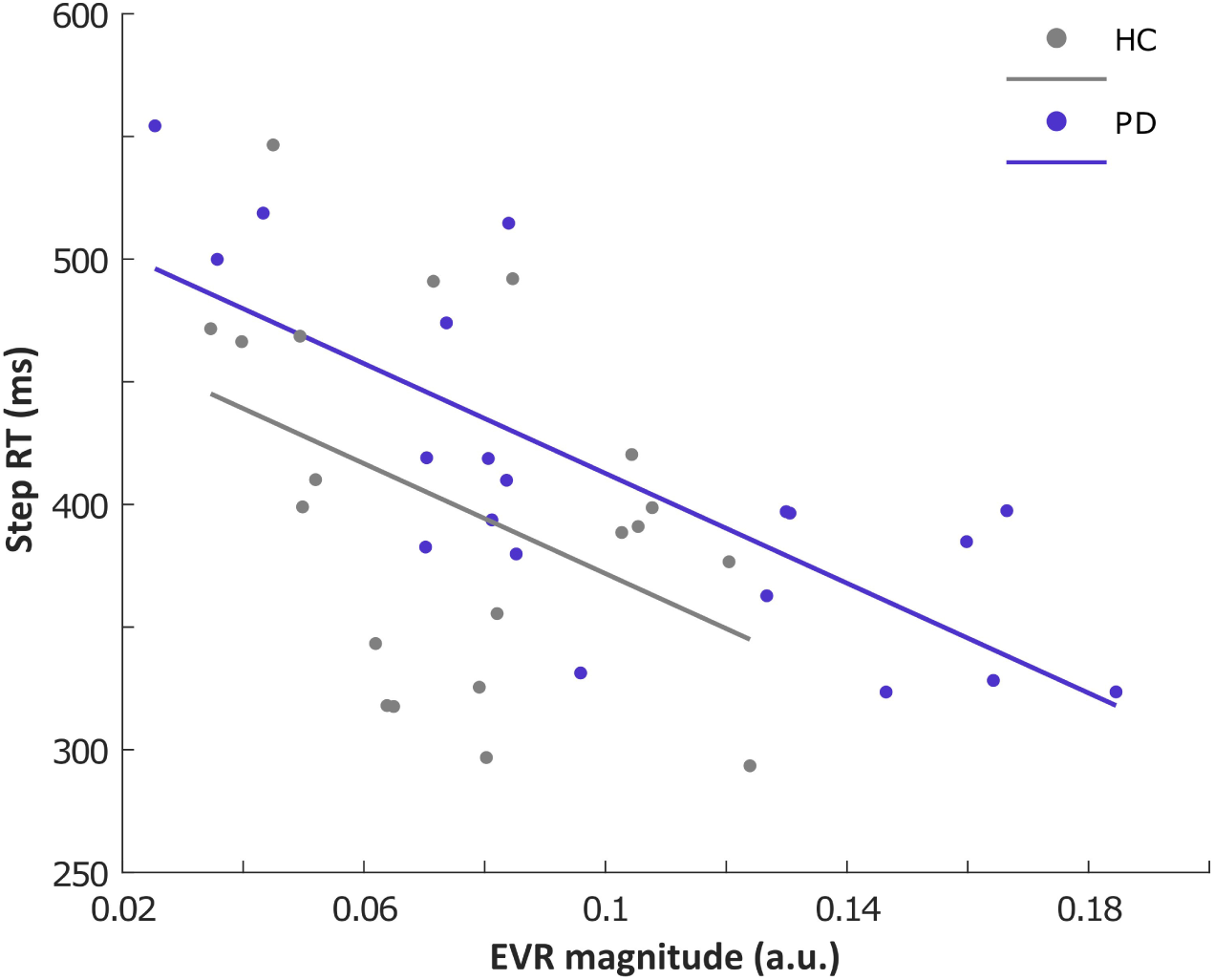
Correlation between response magnitudes in the EVR window and stepping reaction times in healthy elderly participants (purple) and participants with PD (grey) in anterolateral stepping. Note that all participants are displayed here, regardless of whether or not a significant EVR was evoked. The two regression lines to not significantly differ from each other in intercept and slope (p > .05)

As is visible in the two representative participants, there seems to be EVR activity on a subset of slow-RT trials in the anteromedial condition. This was reported previously in our work on a younger healthy cohort (Billen et al., 2023), which we interpreted as a lack of contextual suppression of the EVR the anteromedial condition. In order to test whether the two groups of the current study differed in their ability to contextually suppress the EVR in the anteromedial condition, as in Billen et al. (2023), we split the trials in the anteromedial stepping condition into a fast- and a slow-RT half and separately performing the time-series ROC analyses within the EVR window on the two subsets of trials. Replicating our previous findings, we detected EVRs on the slow half of trials in the majority of the control group (12/20) and the PwPD (13/20; *p* = 1), indicating that both groups had a similar ability to contextually suppress the EVR in the anteromedial condition.

## Do EVR and APA magnitudes correlate with disease progression?

We observed a significant negative correlation (*ρ* = - .59) between response magnitudes within the EVR window and the motor subscale (part III) of the MDS-UPDRS, indicating that PwPD experiencing more severe motor symptoms expressed weaker EVRs (Figure 6A). We further tested whether MDS-UPDRS part III scores differed between participants with and without EVR expression. For both steps towards the right side (T(18) = 0.37, *p* = .71) and steps towards the left side (T(18) = - 1.73, *p* = .11), this effect remained non-significant. We also found no significant correlation between MDS-UPDRS score (part III) and APA magnitudes in the PD group (*ρ* = .26, *p* = .27 Figure 6B).

**Figure 6.**
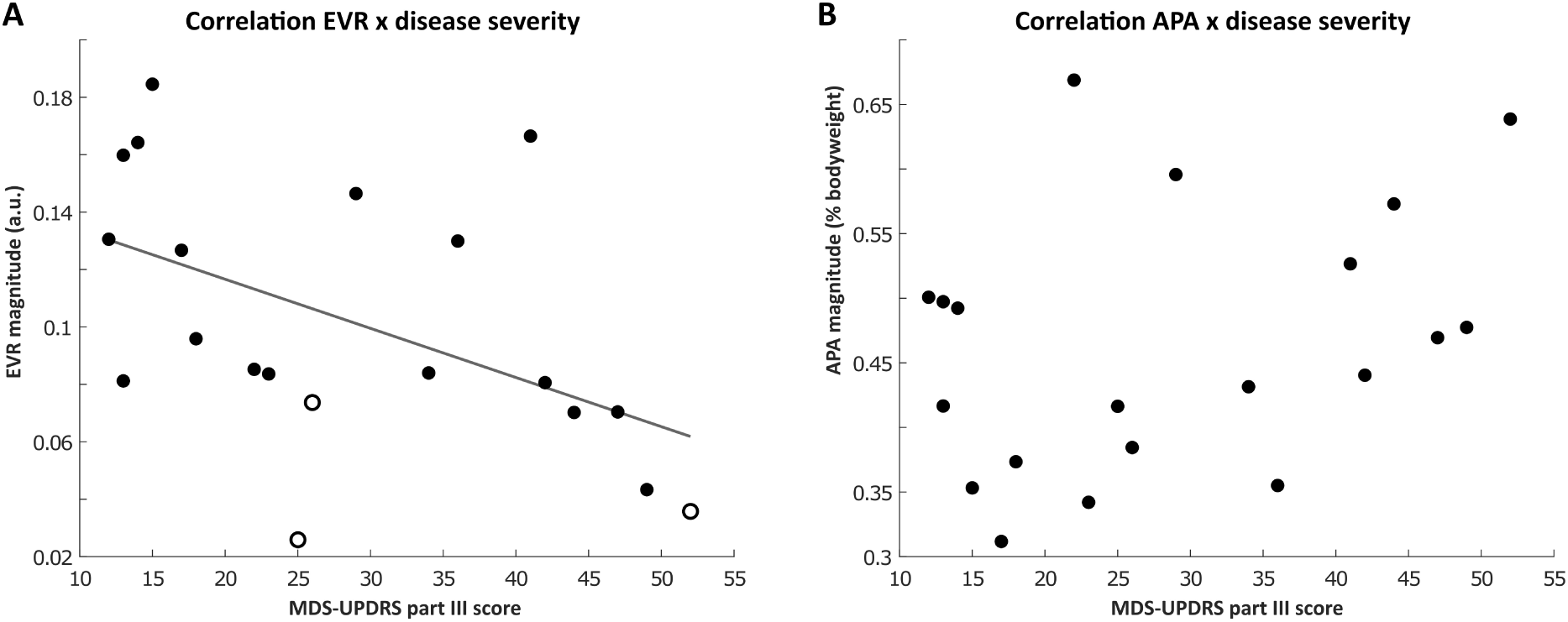
**A:** Correlation between MDS-UPDRS part III scores and EVR magnitudes (averaged across sides) in the PD group during anterolateral stepping. Note that all participants are displayed here, regardless of whether or not a significant EVR was evoked. Participants who did not have EVRs on either side are indicated as black rings. **B:** Scatter plot of MDS-UPDRS part III scores and APA magnitudes in the PD group during anteromedial stepping.

## Discussion

We investigated the relationship between express visuomotor responses (EVRs) and anticipatory postural adjustments (APAs) on the lower limbs of individuals with Parkinson’s Disease (PD) and age-matched healthy controls. From a neuropathological and behavioral point of view, the integrity of the fast visuomotor network has remained relatively unexplored in PD. Our primary objective was to shed light on the intactness of this network by measuring EVRs and APAs in the context of a rapid visually-guided rapid stepping task. EVRs were robustly present in both groups in anterolateral steps. Somewhat surprisingly, PwPD exhibited, on average, stronger EVRs than the control group, but EVR magnitudes decreased with increasing disease severity. In anteromedial stepping, EVRs were largely absent, although most participants from both groups showed EVRs on a subset of trials with slow stepping RTs. While APA magnitudes were decreased in PwPD compared to the control group, subsequent stepping outcomes remained unaffected. Our results demonstrate that the fast visuomotor network that produces EVRs is largely spared in PD, despite concurrent degradation of the circuitry that produces APAs.

## APAs smaller in PD, but subsequent stepping parameters unaffected

In contrast to the extensive research demonstrating severe step initiation and gait impairments in PD (Caetano et al., 2018; Clarke, 2007; Contreras & Grandas, 2012; Palakurthi & Burugupally, 2019), we observed remarkable similarity in stepping-related parameters between PwPD and healthy controls. While APA magnitudes were slightly smaller in the PD group, this did not seem to negatively affect subsequent stepping reaction times, step velocity, duration or size. One potential explanation for the absence of any significant behavioral differences between the two groups could be attributed to the nature of the task itself, which involved rapid step initiation towards highly salient visual stimuli in an emerging target paradigm that is known to enhance both the frequency and magnitude of EVRs. Stepping towards visual stimuli resembles characteristics of classical cueing tasks extensively studied in PD research (Cosentino et al., 2023; Jiang & Norman, 2006; Russo et al., 2022). As internally generated movements involving the basal ganglia posterior putamen are usually impaired in PD due to dopaminergic depletion, external cueing has been proposed to engage alternative pathways involving corticostriatal loops, thereby bypassing some of the more strongly affected areas (Cosentino et al., 2023; Tosserams et al., 2022). In the current study, a similar mechanism might be at play, whereby the salient visual stimulus in combination with the goal of stepping toward the stimulus may have circumvented some of the more severely affected neural circuits, thereby masking some of the deficits experienced in daily life. Interestingly, APAs were indeed smaller in the PD group during anteromedial stepping, while the subsequent stepping reaction times remained unaffected. These findings may point towards an altered speed-accuracy trade-off in the PwPD compared to the control participants. PwPD generally seemed to prioritize speed at the cost of accuracy, underlined by slightly increased error rates, similar to previous findings on interceptive movements in the upper limb (Fooken et al., 2022), and in an anti-reach task (Gilchrist et al., 2024).These findings may hint at PwPD tending to push the boundaries of a safe step more than the healthy controls did, by lifting the foot despite a smaller and potentially insufficiently strong APA, possibly leading to smaller stability margins upon foot landing.

## EVR expression largely spared in people with PD

Our results revealed that EVRs were robustly expressed in the majority of PwPD (17/20 participants) and in the healthy control group (16/20 participants). While EVR latencies did not differ between groups, EVR magnitudes were, on average, even larger in the PD group compared to the HC group. These findings suggest that the EVR network is relatively spared in PD, which is in line with recent findings from a study on upper-limb EVR expression in PD (Gilchrist et al., 2024). Another important aspect of EVR expression is adequate context-dependent suppression when the postural demands are high. In this context, EVRs are counterproductive on anteromedial steps, as they propel the center of mass forward. The majority of the participants from both groups expressed EVRs on the slow half of trials in anteromedial stepping, suggesting that the (presumable) top-down suppression of EVRs occasionally lapsed on a subset of trials in most participants, which required larger and longer lasting APAs in the stepping leg, and consequently longer stepping reaction times. These findings are in line with the observed expression of slow-half anteromedial EVRs in a majority of younger participants from a previous study (Billen et al., 2023), suggesting that, while not perfect, adequate EVR suppression in the PD group was not additionally affected. It is important to note that due to the blocked design of the current task, participants were able to proactively prepare for the expected postural demands of the upcoming step. It would be of interest to investigate the contextual interaction between EVRs and postural control in an intermixed design, whereby postural demands are manipulated on a trial-to-trial basis. In such an unpredictable context, behavioral and EVR-related differences between PwPD and the HC control group may start to emerge, as recent evidence suggests that increased task demands in PwPD hampers their ability to contextually suppress or govern the EVR in the upper limb (Gilchrist et al., 2024).

Despite the present and other recent results suggesting general sparing of the fast visuomotor network in PD, we found that EVR magnitude progressively declined as disease severity increased, albeit without significant differences in MDS-UPDRS part III scores between participants with PD who expressed EVRs and those who did not. Intriguingly, the average EVR magnitude of the subgroup of mildly affected PwPD (MDS-UPDRS part III scores of 12-18) was on average higher (∼ 0.12 a.u.) than the mean EVR magnitude of the healthy control group (0.08 a.u.), suggesting that EVRs may be upregulated in the early stages of the disease, potentially as part of the greater speed-accuracy tradeoff as discussed above and as previously reported (Fooken et al., 2022; Gilchrist et al., 2024). Simultaneously, upregulation of the EVRs may be a mechanism to compensate for some of the early motor deficits that originate in more severely affected areas. However, as the PD symptoms progress and more neural circuits become affected, the fast visuomotor system may start to be afflicted as well, as reflected in decreased EVR magnitudes. The exact neuropathological mechanisms of EVR degradation remain to be established. Even though EVRs are likely generated in superior colliculus, top-down input from various other areas can modulate the network and either upregulate or downregulate EVRs depending on the context-dependent task demands (Contemori et al., 2021b, 2023; Gu et al., 2016). The observation that EVR output weakens with increasing disease severity may thus be due to the SC itself becoming affected as the disease progresses and/or to degradation of the “priming” inputs to SC, such as basal ganglia or higher-order cortical areas. In the current study, we cannot discriminate between these two possibilities, so this remains an essential question for future research.

## Healthy elderly participants performed worse compared to healthy young participants

Compared to a cohort of younger participants from a previous study (*M_Age_* = 23.3 years; Billen et al., 2023), the current cohort of healthy elderly participants exhibited EVRs at a lower prevalence (80% of participants here, compared to 100% in previous study), and longer latencies (∼120 ms in the elderly participants here, compared to ∼108 ms in the previous study). The elderly also initiated steps more slowly (average RT in anterolateral stepping of 399ms compared to 314ms in previous study). Several factors could contribute to these disparities. Firstly, it is plausible that age-related physiological changes (e.g. weaker muscles, cognitive decline, slower sensory integration; for review, see Osoba et al., 2019) may have rendered older participants physically less capable of performing as robustly as the younger participants, similar to previous studies reporting age-related changes in gait- and balance-related behavior (Boyer et al., 2023; Dewolf et al., 2021; Reimann et al., 2020; Zhang et al., 2021). Additionally, psychological factors might come into play, as with age, individuals may prioritize safe stepping due to the potentially greater consequences of falls. This heightened focus on maintaining balance could explain why EVRs are generally weaker and less prevalent in the elderly, which contributes to the longer stepping reaction times. However, this cautious approach, if taken to extremes, might lead to inflexibility in adapting to sudden environmental changes, which could increase the risk of falls (Weerdesteyn et al., 2004). Understanding these dynamics is important for improving our understanding of motor behavior across different age groups and may inform strategies for fall prevention in elderly populations.

## Limitations

It should be noted that PwPD were not asked to deviate from their usual medication schedule in the current experiment, meaning that all participants generally performed the tasks in the ON state. This approach contrasts with previous studies on upper limb EVRs (Gilchrist et al., 2024) and other research on online corrections (Desmurget et al., 2004), where participants performed the experiment while in the OFF state. Of note, Merritt et al. (2017) investigated the effect of dopaminergic therapy on fast online corrections, reporting only minor effects. Because not all movement symptoms are affected equally by dopaminergic therapy, it is plausible that the rapid, stimulus-driven and goal-directed movements used in these types of studies are less sensitive to dopaminergic medication. In the context of stepping, it would be valuable for future research to investigate the effect of dopaminergic medication on EVR expression and postural control, as this could provide deeper insights into the role of medication in these processes.

## Conclusion

Here we provide compelling evidence that, at least in the context of a rapid, goal-directed stepping task used here, the participants with PD were able to show unaffected stepping behavior. This is remarkable, given the deficits experienced in daily life and the large disease severity range present in the current study. We speculate that the nature of our stepping task resembled a visual cueing task, which has previously been shown to help in overcoming some of the deficits experienced in daily life. Future research should focus on the specific context-dependent variables that promote the overcoming of impaired stepping behavior, potentially in even more severely affected participants, such as PwPD that experience freezing of gait.

We also demonstrate relatively spared EVRs in people with PD during step initiation. Especially in the early stages of the disease, EVR output may in fact be upregulated, potentially to compensate for more affected aspects of the motor output. Together with evidence for EVR expression on upper limb muscles, this further provides insight into the neuropathological mechanisms underlying the fast visuomotor network. Future studies may investigate the role of this network in more complex contexts that more closely resemble the unpredictability of our dynamic environment.

## Author Contributions

*Lucas Billen:* Conceptualization, Methodology, Software, Formal analysis, Investigation, Data curation, Writing – Original Draft, Review & Editing, Visualization; *Brian Corneil:* Conceptualization, Methodology, Software, Resources, Writing - Review & Editing, Supervision, Funding acquisition; *Vivian Weerdesteyn:* Conceptualization, Methodology, Resources, Writing - Review & Editing, Supervision, Project administration, Funding acquisition

## Acknowledgements

Thank you to Ilse Giesbers for helping during the early stages of the project.

## Data availability statement

The authors confirm that the data supporting the findings of this study are available within the article and its supplementary materials.

